# Importance of transcript variants in transcriptome analyses

**DOI:** 10.1101/2024.07.11.603122

**Authors:** Kevin Vo, Ryan Mohamadi, Yashica Sharma, Amelia Mohamadi, Patrick E. Fields, M. A. Karim Rumi

## Abstract

RNA sequencing (RNA-Seq) has become a widely adopted genome-wide technique for investigating gene expression patterns. However, conventional RNA-Seq analyses typically rely on gene expression (GE) values that aggregate all the transcripts produced by a gene under a single identifier, overlooking the complexity of transcript variants arising from different transcription start sites and alternative splicing events. In this study, we explored the implications of neglecting transcript variants in RNA-Seq analyses. Among the 1334 transcription factor (TF) genes expressed in mouse embryonic stem (ES) or trophoblast stem (TS) cells, 652 were reported to be differentially expressed in TS cells based on GE values (365 upregulated and 287 downregulated, ≥2-fold, FDR *p*-value ≤0.05). Intriguingly, differential gene expression analysis revealed that of the 365 upregulated genes, 883 transcript variants were expressed, with only 174 (<20%) variants exhibiting upregulation based on transcript expression (TE) values. The remaining 709 (>80%) variants were either down-regulated or showed no significant change in expression analysis. Similarly, the 287 genes reported to be downregulated expressed 856 transcript variants, with only 153 (<20%) downregulated variants and 703 (>82%) variants that were upregulated or showed no significant changes. Additionally, the 682 TF genes that did not show significant changes between ES and TS cells (GE values < 2-fold changes and/or FDR p-values >0.05) expressed 2215 transcript variants, which included 477 (>21%) that were differentially expressed (276 upregulated and 201 downregulated, ≥2-fold, FDR p-value ≤0.05). Notably, a particular gene does not express just one protein; rather its transcript variants encode multiple proteins with distinct functional domains, including non-coding regulatory RNAs. Our findings underscore the critical necessity of considering transcript variants in RNA-Seq analyses. Doing so may enable a more precise understanding of the intricate functional and regulatory landscape of genes; ignoring the variants may result in an erroneous interpretation.

**Graphic Abstract:** 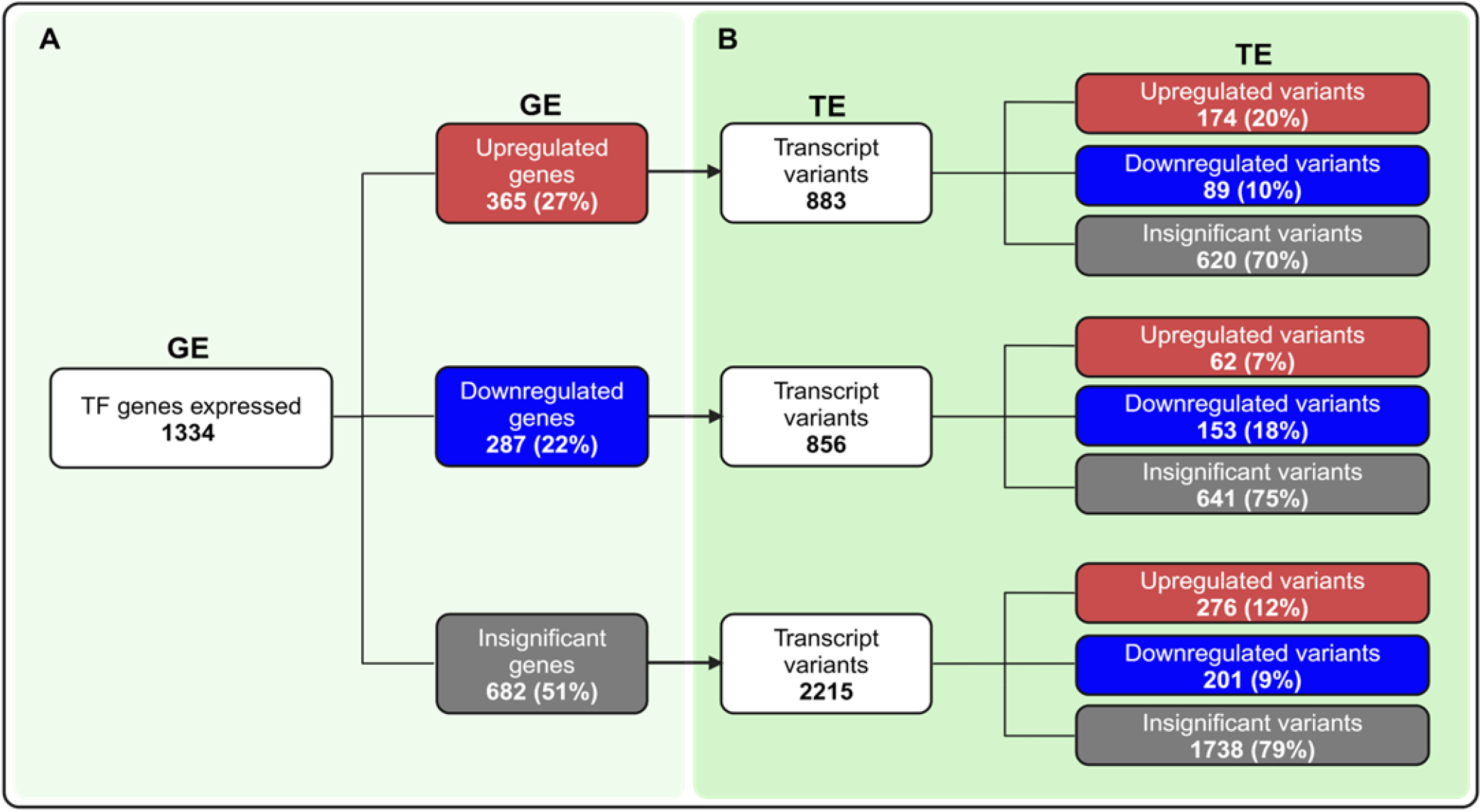

Differential expression of transcription factors (TFs) between mouse embryonic stem (ES) cells and trophoblast stem (TS) cells. This graphic presentation clearly demonstrates the importance of including transcript variants during RNA sequencing (RNA-Seq) analyses. Panel **A** represents the conventional differential gene expression analysis approach after RNA-Seq, where all transcript reads are taken under a single gene name. Panel **B** takes differential gene expression analysis one step further by examining all the transcript variants that were previously hidden under the main gene name. Our results indicate that exclusive gene expression (GE) analysis inaccurately defines over 80% of the transcript expression (TE). Without analyses of all the transcript variants’ reads, we fail to uncover the functional importance of the variants and the regulation of their expression. Both GE and TE values are expressed as transcript per million (TPM). Data analyses were performed by using CLC Genomics Workbench.

## 1. Introduction

Understanding gene expression at the cellular level is crucial for unraveling cell-type-specific functions, identifying biomarkers, and pinpointing genes or pathways for targeted molecular interventions^1^. RNA sequencing (RNA-Seq) has emerged as a powerful tool for comprehensive transcriptome analysis, enabling the identification of lineage-specific gene expression patterns^2-4^. Integrating RNA-Seq with techniques such as ATAC-Seq, ChIP-Seq, Cut & Run, Ribo-Seq, and methyl-Seq has provided insights into the intricate interplay between epigenomic modifications and transcriptional regulation^5, 6^. Moreover, a detailed examination of the transcriptome offers a window into the gene regulatory mechanisms within a distinct cell type^7^.

A single gene does not express a single mRNA to encode a single protein^8^. Commonly, multiple mRNAs that are transcribed encode different proteins or noncoding RNAs^8^. Alternative transcription start sites (ATSS) can result in the expression of more than one transcript from a single gene^9^. Alternative transcription start sites occur due to alternative proximal promoter use as well as the availability of alternative transcriptional regulators in a particular cell type^10, 11^. However, alternative splicing is a common mechanism underlying the generation of multiple mRNA variants from a single initial transcript^12^. RNA editing may further expand the repertoire of transcript variants^13^. Although the transcript variants are translated into peptides using the same open reading frame, they can encode various proteins with different lengths and functional domains^14^. Some alternative transcripts also do not encode any proteins that may act as long noncoding RNAs or other regulatory RNAs^15^. Thus, alternative transcripts, some of which encode non-coding RNAs, play pivotal roles in lineage-specific divergent cellular functions.

Despite the functional significance of transcript variants, conventional gene expression analyses typically overlook the diversity of transcripts. Current RNA-Seq methodologies often quantify gene expression (GE) values in Reads Per Kilobase Million (RPKM) or Transcripts Per Million (TPM), aggregating all transcript counts under a single mRNA identifier without distinguishing between full-length transcripts and their variants, irrespective of their protein-coding potential^16, 17^. However, RNA-seq analyses can generate quantitative data regarding transcript variants’ expressions (TE values). While transcript expression (TE) values can be concurrently calculated, GE values predominantly drive the identification of differentially expressed genes across experimental conditions, largely due to analytical complexities and validation challenges hindering the widespread adoption of TE analyses^18^.

It is physiologically inaccurate to consider the expression of a single transcript from a specific gene for differential expression analyses. In addition, it can also be misleading to conclude that similar trends of expression occur for all the transcript variants expressed from a single gene. Therefore, we have evaluated the limitations of GE-based RNA-Seq analyses that fail to consider the TE values of transcript variants. Our results indicate that GE-based RNA-seq analyses incorrectly represented over 80% of the TE-based analyses.

## 2. Materials and Methods

### 2.1 Experimental Model

We have used RNA sequencing data of two early embryonic stem cell lines, embryonic stem (ES) cells, and trophoblast stem (TS) cells, and focused on transcription factors (TFs). Differential expression of lineage-specific TFs are characteristic determinants of ES and TS cell lineages. Ectopic expression of selective lineage-specific TFs can reprogram somatic cells into ES or TS cells ^19, 20^ (**Figure 1A**). Moreover, TFs are appropriate for defining the role of transcript variants due to their well-defined functional domains (DNA binding domain-DBD, transactivation domain-TAD, and signaling sensing domain-SSD)^21^ (**Figure 1B**). Thus, transcript variants of the same TF gene may encode proteins carrying different DBD, TAD, or SSD that can be easily determined. We have systematically analyzed the differential expression of TFs between the ES and TS cells to understand the limitations of transcriptome analyses without considering the transcript variants.

**Figure 1.**
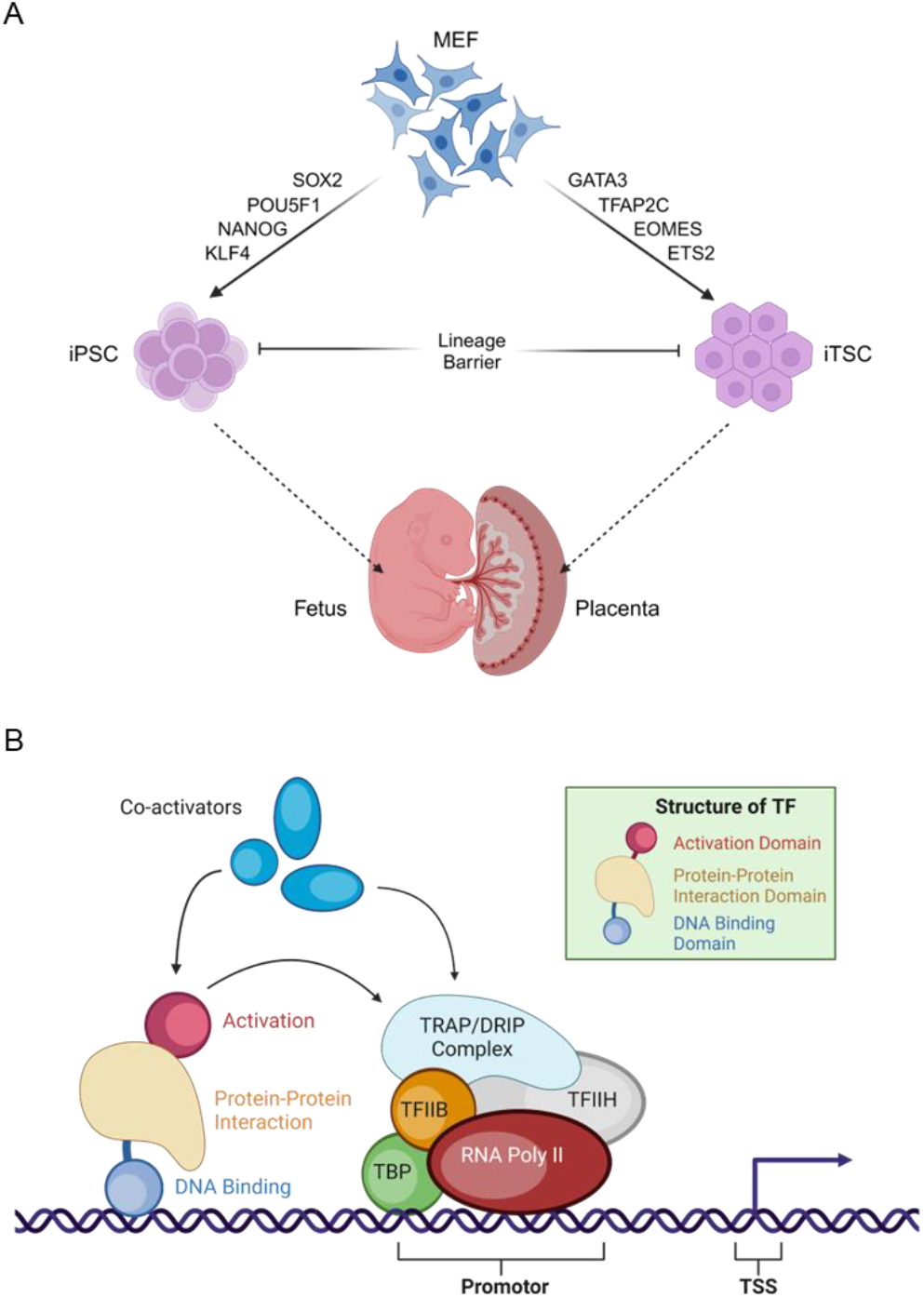
Transcription factors and early embryonic stem cells. The schematics explain the justification for choosing TFs for this study and using RNA-seq data of mouse ES cells and TS cells (A). Transcript variants encoding distinct domains are easily understandable in TFs, and the role of TFs in determining cell fate is well known (B). Arrow indicates start of gene transcription. MEF, mouse embryonic fibroblast; iPSC, induced pluripotent stem cells; iTSC, induced trophoblast stem cells; TF, transcription factor; RNA Pol II, RNA polymerase II; TSS, transcription start site.

### 2.2 RNA Sequencing Data

In this study, we included RNA-seq data from mouse ES cells (n=3 libraries) and mouse TS cells (n = 3 libraries). Mouse TS cell data were generated in our laboratory and have been submitted to the Sequencing Read Archive (PRJNA1131096; SRA, NCBI). The ES cell data were downloaded from the NCBI’s Gene Expression Omnibus (GEO) (SRR24044798, SRR24044809, SRR24044810) ^22^.

Mouse TS cells were maintained in feeder-free stem conditions^23^. TS cells were cultured for 48 hours, and total RNA was extracted using TRI reagent (Millipore-Sigma, St. Louis, MO, USA). 500 ng of total RNA from each sample (RIN value >9) was used for the sequencing library preparation using a TruSeq Stranded mRNA kit (Illumina, San Diego, CA, USA). The cDNA libraries were evaluated for quality at the KUMC Genomics Core and sequenced on an Illumina HiSeq X sequencer at Novogene Corporation (Sacramento, CA, USA).

### 2.3 RNA Sequencing Analysis

RNA-Seq data were analyzed using CLC Genomics Workbench (Qiagen Bioinformatics). All clean reads were obtained by removing low-quality reads and trimming the adapter sequences. The high-quality reads were aligned to the Mus musculus reference genome (GRCm39), gene (GRCm39.111_Gene), and mRNA sequences (GRCm39.111_mRNA) using the default parameters: (a) maximum number of allowable mismatches was 2 and (b) minimum length and similarity fraction was set at 0.8, and (c) a minimum number of hits per read was 10. The expression values of individual genes (GE) or transcript variants (TE) in ES and TS cells were determined as described in our previous publications ^24-26^. Expression values were measured in TPM.

Expression of 16,052 to 16,510 genes was detected in the ES or TS cell-derived RNA-seq samples. We selectively analyzed the 1,374 mouse TFs that were curated by the Gifford lab (https://cgs.csail.mit.edu/ReprogrammingRecovery/mouse_tf_list.html) from a list of human TFs ^27^. New tracks containing only the TFs were generated from each RNA-Seq data file containing GE or TE values, which were used in subsequent analyses. The threshold p-values were determined according to the false discovery rate (FDR) to identify the differentially expressed genes or transcript variants between ES and TS cells. A gene or a transcript variant was considered differentially expressed if the absolute fold change was ≥2 and the FDR p-value was ≤ 0.05 ^24-26^.

### 2.4 Analysis of the Transcript Variants

We analyzed the differential expression of genes using the RNA-seq files containing GE values. The differentially expressed genes were divided into 3 groups: upregulated (≥2-fold changes and FDR *p* ≤0.05), downregulated (≤ -2-fold changes and FDR *p* ≤0.05), and insignificant (either ≤ absolute 2-fold changes and/or FDR *p* ≥ 0.05). The transcript variants encoded by the upregulated, downregulated, or insignificant group of genes were further analyzed to identify the differentially expressed ones between mouse ES and TS cells. Differential expressions of the transcript variants were analyzed using the RNA-seq files containing TE values. These analyses identified the differentially upregulated (≥2-fold changes and FDR *p* ≤0.05), downregulated (≤ -2-fold changes and FDR *p* ≤0.05), and insignificant (either ≤ 2-fold changes or FDR *p* ≥ 0.05) group of transcript variants.

### 2.5 Statistical Analyses

For RNA Seq, each study group contained three library samples. In CLC Genomics Workbench, the ‘differential expression for RNA-Seq tool’ performs some multi-factorial statistics on a set of expression tracks based on a negative binomial generalized linear model (GLM). The final GLM-fit and dispersion estimate calculates the total likelihood of the model given the data and the uncertainty of each fitted coefficient^28^. Two statistical tests-the Wald and the Likelihood Ratio tests-use one of these values. The Across groups (ANOVA-like) comparison uses the Likelihood Ratio test.

## 3. Results

### 3.1 Lineage-specific Expression of Transcription Factors

Expression of the TFs in mouse ES cells and TS cells was analyzed using RNA-seq data. Differential expression of TF genes between ES and TS cells was evident in the heat map (**Figure 2A**). Based on the Pearson correlation matrix of the TFs, there was a high positive relation among the 3 ES samples, or the 3 TS samples (**Figure 2A**).

**Figure 2.**
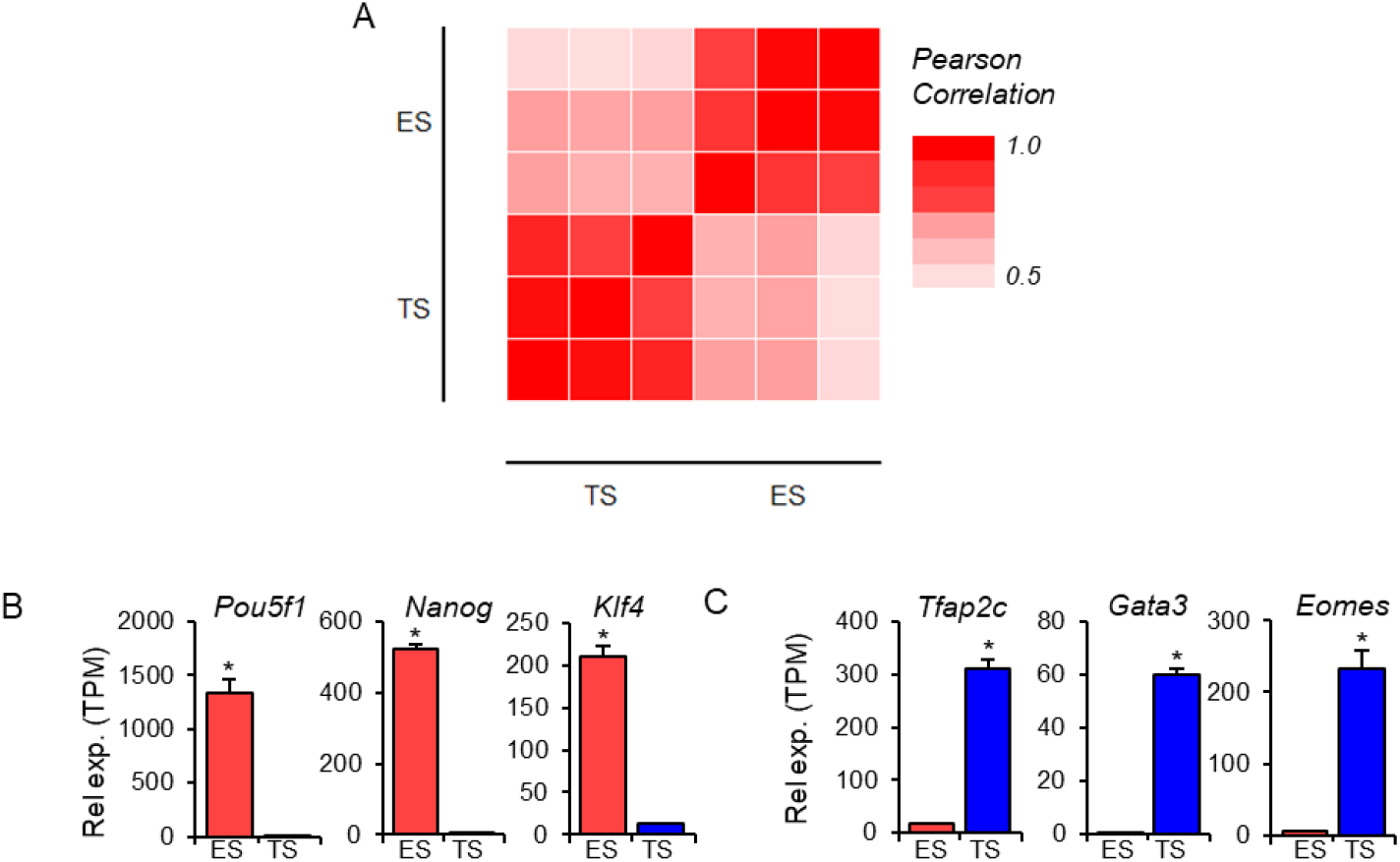
Quality and validity of the RNA-seq data obtained from mouse ES cells and TS cells. (**A**) A matrix shows the Pearson correlation of 1365 transcription factors (TFs) expressed in 3 ES and 3 TS samples. ES-specific abundant expression of characteristic TFs (*Pou5f1, Nanog*, and Klf4) that differentiate them from TS cells (B) and TS-specific abundant expression of TFs (Tfap2c, Gata3, and Eomes) that distinguish those from ES cells (C) indicate the RNA-seq data quality and validity. Data represent mean TPM±SE, * indicates *p* < 0.05. Rel exp., relative expression; TPM, transcript per million.

The validity of ES and TS lineage identity was confirmed based on the expression of stem cell markers (*Pou5f1, Nanog*, and *Klf4* for ES; *Tfap2C, Gata3*, and *Eomes* for TS) ^29-31^. High levels of *Pou5f1, Nanog*, and *Klf4* were expressed in mouse ES cells but were very low in TS cells (**Figure 2B**). In contrast, *Tfap2c, Gata3*, and *Eomes* expressions were very high in mouse TS cells but low in ES cells (**Figure 2C**).

### 3.2 Differential Expression of the Transcription Factor Genes and Transcript Variants

Of the 1,374 TFs, 1,334 were expressed in mouse ES or TS cells. TS cells showed differential expression of 652 TF genes compared to ES cells (365 upregulated and 287 down-regulated; ≥ 2-fold changes, FDR *p*-value <0.05) (**Figure 3 A-C**). The differential expressions of the GE values in TS cells are evident in heatmaps (**Figure 3A**), volcano plots (**Figure 3B**), and bar graphs (**Figure 3C**). The 1,334 TF genes expressed 3,954 transcript variants in mouse TS or ES cells (**Figure 3 D-F**). A total of 1,739 of the 3,954 transcript variants were differentially expressed in TS cells (883 upregulated and 856 downregulated; ≥ 2-fold changes, FDR *p*-value <0.05) (**Figure 3 D-F**). The differential expressions of the TE values in TS cells are shown in heatmaps (**Figure 3D**), volcano plots (**Figure 3E**), and bar graphs (**Figure 3F**).

**Figure 3.**
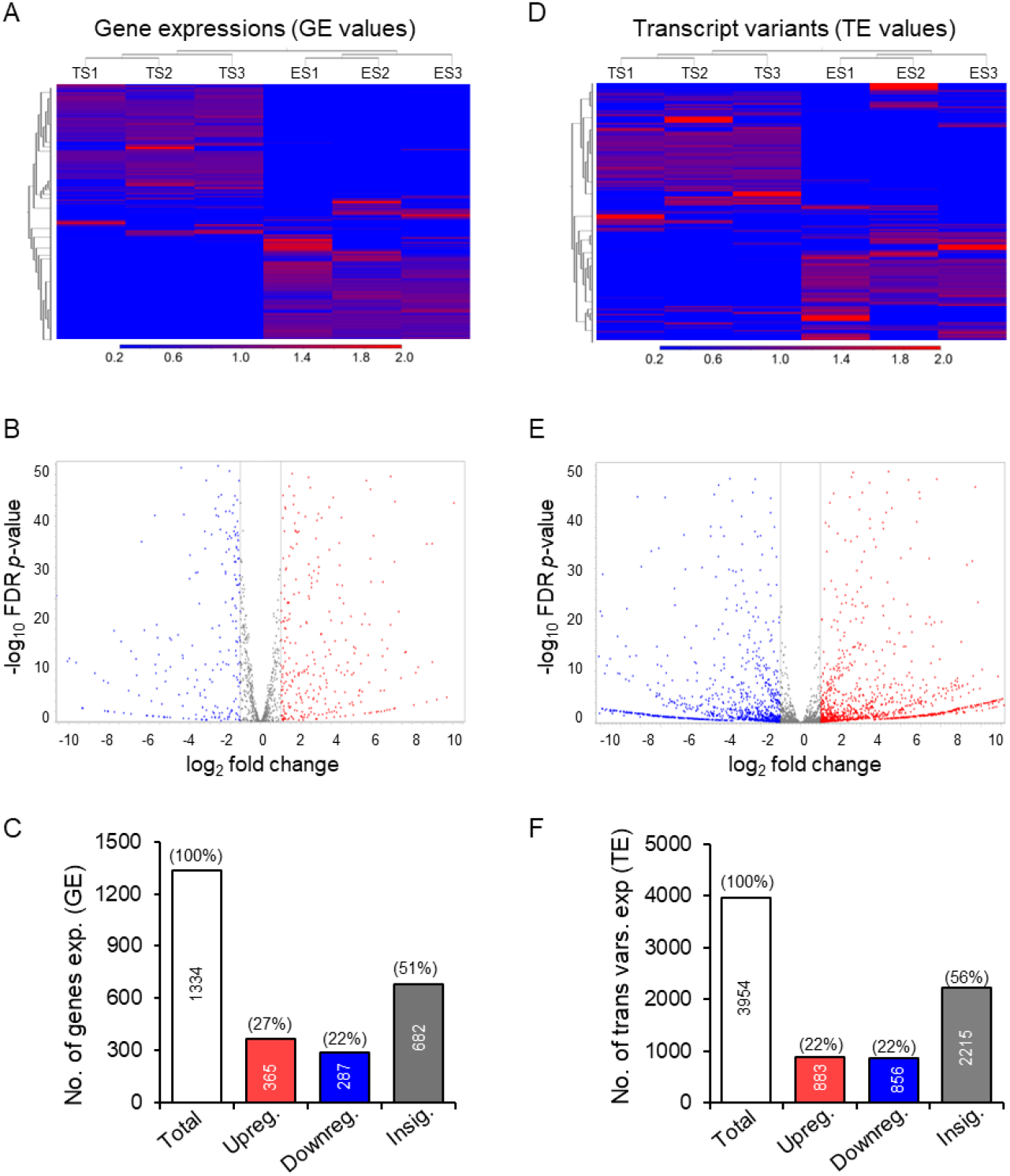
Differential expression of genes and transcript variants in mouse TS cells compared to ES cells. Heat maps, Volcano plots, and bar graphs show that ∼49% of the genes were differentially expressed (27% upregulated and 22% downregulated) in TS cells (**A-C**). Similarly, 44% of the transcript variants encoded by the TF genes were differentially expressed (22% upregulated and 22% downregulated) in TS cells. A 5% reduction in upregulated transcript variants was associated with increased variants in the insignificant group.

### 3.3 Discrepancy between Gene Expression and Transcript Variants

Despite the overall similar differential expression of genes (based on GE values) and transcript variants (based on TE values) (**Figures 3 C** and **F**), further analyses revealed a remarkable discrepancy between GE and TE-based analyses **(Figures 4** and **5**). The 365 upregulated genes in TS cells expressed 883 transcript variants. Of those 883 transcript variants, only 174 showed significant upregulation (≥ 2-fold upregulation, FDR p-values ≤ 0.05). The remaining 89 transcript variants were significantly downregulated (≥ 2-fold downregulation, FDR p-values ≤ 0.05), and 620 showed insignificant differences based on TE values in TS cells (**Figure 4 A, D**). The 287 downregulated genes expressed 856 transcript variants, of which only 153 were significantly downregulated (≥ 2-fold downregulation, FDR *p*-values ≤ 0.05). The remaining 62 transcript variants were upregulated (≥ 2-fold upregulation, FDR *p*-values ≤ 0.05), and 641 showed insignificant differences based on TE values (**Figure 4 B, D**). The 682 genes that showed no significant differential expression based on GE values (absolute fold changed <2 or FDR p-values > 0.05) contained 2215 transcript variants. Of those, 276 transcripts showed significant upregulation (≥ 2-fold upregulation, FDR p-values ≤ 0.05), and 201 showed significant downregulation (≥ 2-fold downregulation, FDR p-values ≤ 0.05) (**Figure 4 C, D**).

**Figure 4.**
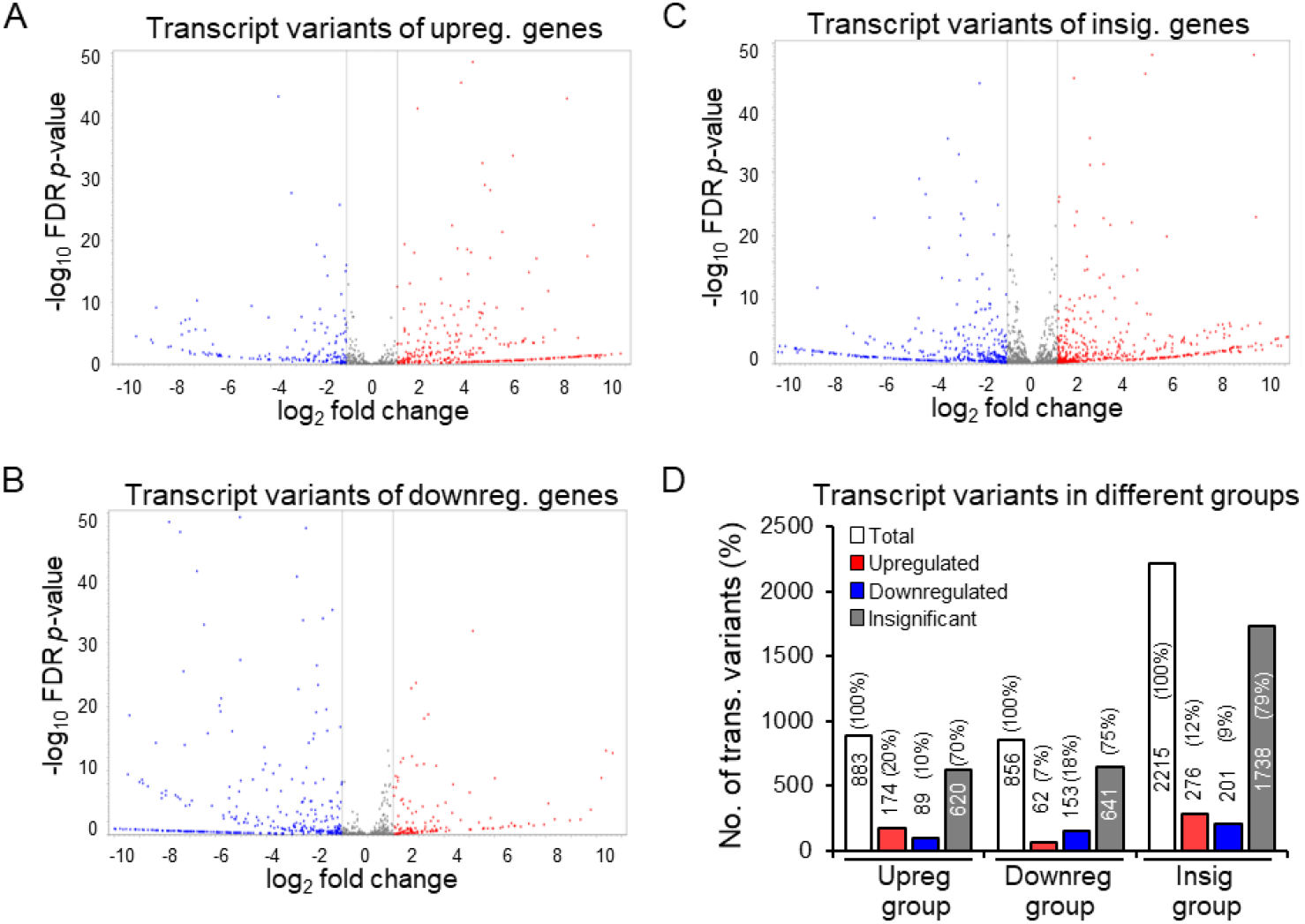
Discordant differential expression of the transcript variants expressed in mouse TS cells. Volcano plots show the differential expression of transcript variants corresponding to upregulated (A), downregulated (B), and insignificant genes (C). ∼80% of transcript variants of the upregulated genes were either downregulated or insignificant (D). Similarly, ∼82% of transcript variants of the downregulated genes were either upregulated or insignificant (D). In addition, ∼21% of transcript variants of the insignificant genes were differentially expressed (D).

### 3.4 Increased Discrepancy among the Low Abundant Transcript Variants

We further analyzed the transcript variants among the differentially expressed genes according to their abundance in mouse TS or ES cells (**Figure 5 A-H**). The low abundant transcript variants (TPM <5 TE values) showed greater discrepancy compared to the moderately high abundant transcripts (TPM ≥5 TE values) (**Figure 5 A-H**). The 365 upregulated genes expressed 883 transcript variants, 666 were low abundant and 217 were moderate to high abundant (**Figure 5 G, H**). The 287 downregulated genes expressed 856 transcript variants, 578 were low abundant and 278 were high abundant (**Figure 5 G, H**). The 682 insignificant genes included 2215 transcript variants, 1476 were low abundant (TPM <5 TE values) and 739 were high abundant (TPM ≥ 5 TE values) (**Figure 5 G, H**).

**Figure 5.**
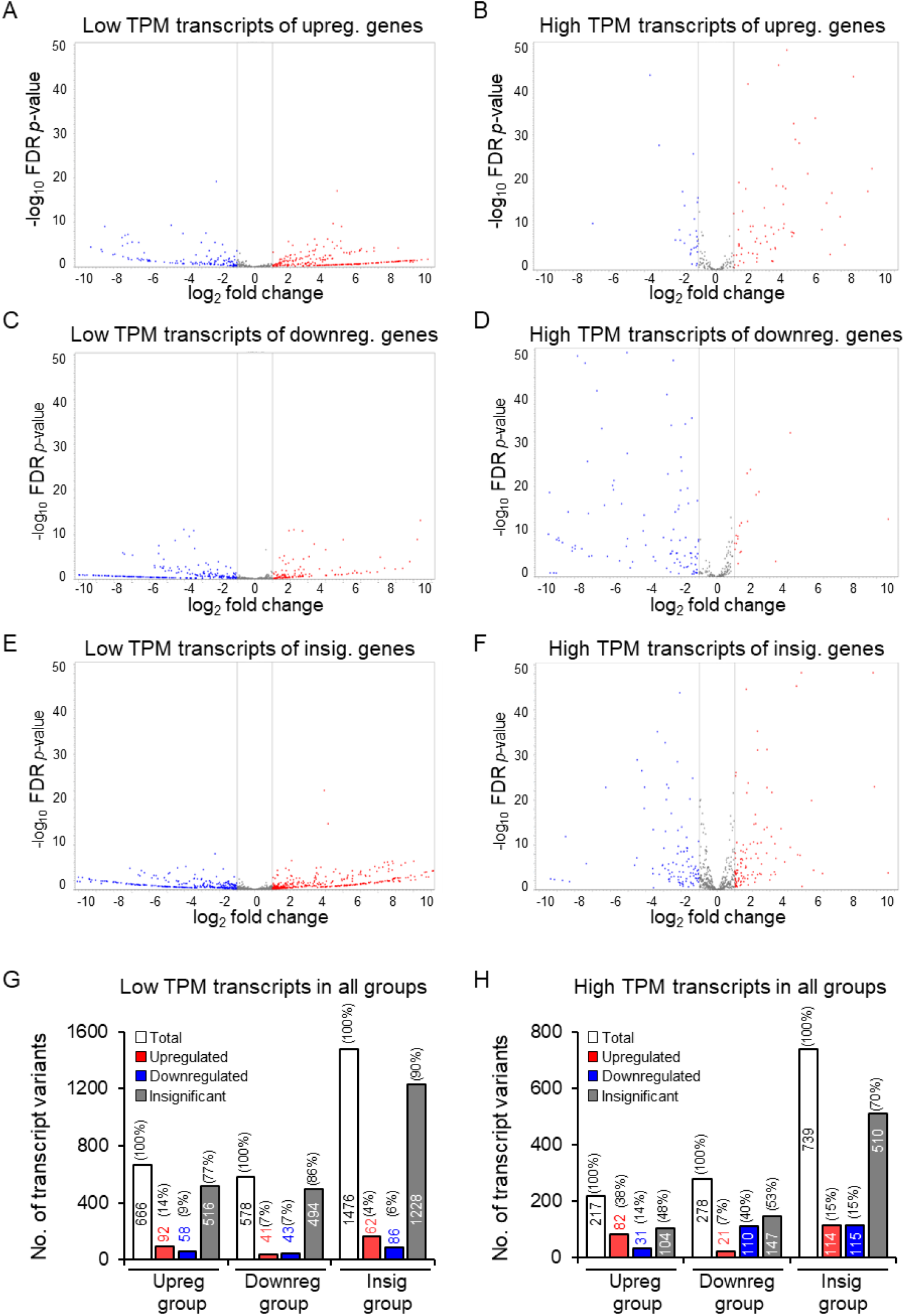
Differential expression of the low copy and high copy number transcript variants expressed in mouse ES or TS cells. Volcano plots showed discordant results among the low copy (< 5 TPM) number (**A, C**, and **E**) as well as the high copy number (**B, D**, and **F**) transcript variants expressed by upregulated (**A**, B), downregulated (**C, D**) or insignificant (E, F) genes. While the low copy number transcripts of the upregulated genes showed discordant results in 86%, it was only 62% among the high copy number transcripts (**G, H**). Similarly, 93% of the low-copy number transcripts encoded by downregulated genes were discordant, and only 60% of the high-copy number genes were discordant (G, H). Remarkably only 10% of transcript variants of insignificant genes were differentially expressed, whereas it was 30% for the high copy number genes.

We observed that ∼86% of the low copy number transcripts of upregulated genes showed discordant results (**Figure 5 A, G, H**). In contrast, ∼62% of the high copy number transcripts of upregulated genes showed discrepant results (**Figure 5 B, G, H**). Similarly, 93% of the low copy number transcripts and 60% of the high copy number transcripts that were expressed by the downregulated genes were discrepant. (**Figure 5 C, D, G, H**). Among the low abundant transcript variants expressed by the insignificant genes, only ∼10% showed differential expression, whereas it was ∼30% among the high copy number transcript variants (**Figure 5 E-H**).

### 3.5 The Basis of Discrepancy between Gene Expression and Transcript Variants Analyses

Our next step of investigations was directed towards elucidating the molecular basis of the discrepancy between analyses of gene expressions and transcript variants. In this analysis, we included two from each group of the upregulated, downregulated, or insignificant genes that demonstrated a discrepancy between their GE-based and TE-based analyses (as identified in Section 3.4) (**Figure 5**). Here, we have analyzed the transcript variants of Kmt2a and Hmg20b from the GE-based upregulated group (Figure 6), Gtf2i and Rbpj from the downregulated group (Figure 7), and *Atf2* and *E2f3* genes from the insignificant group (**Figure 8**). We identified that the downregulation of a transcript variant can be masked by relatively higher upregulation of another transcript variant while they were encoding proteins with different functional domains (**Figure 6 A, B**).

**Figure 6.**
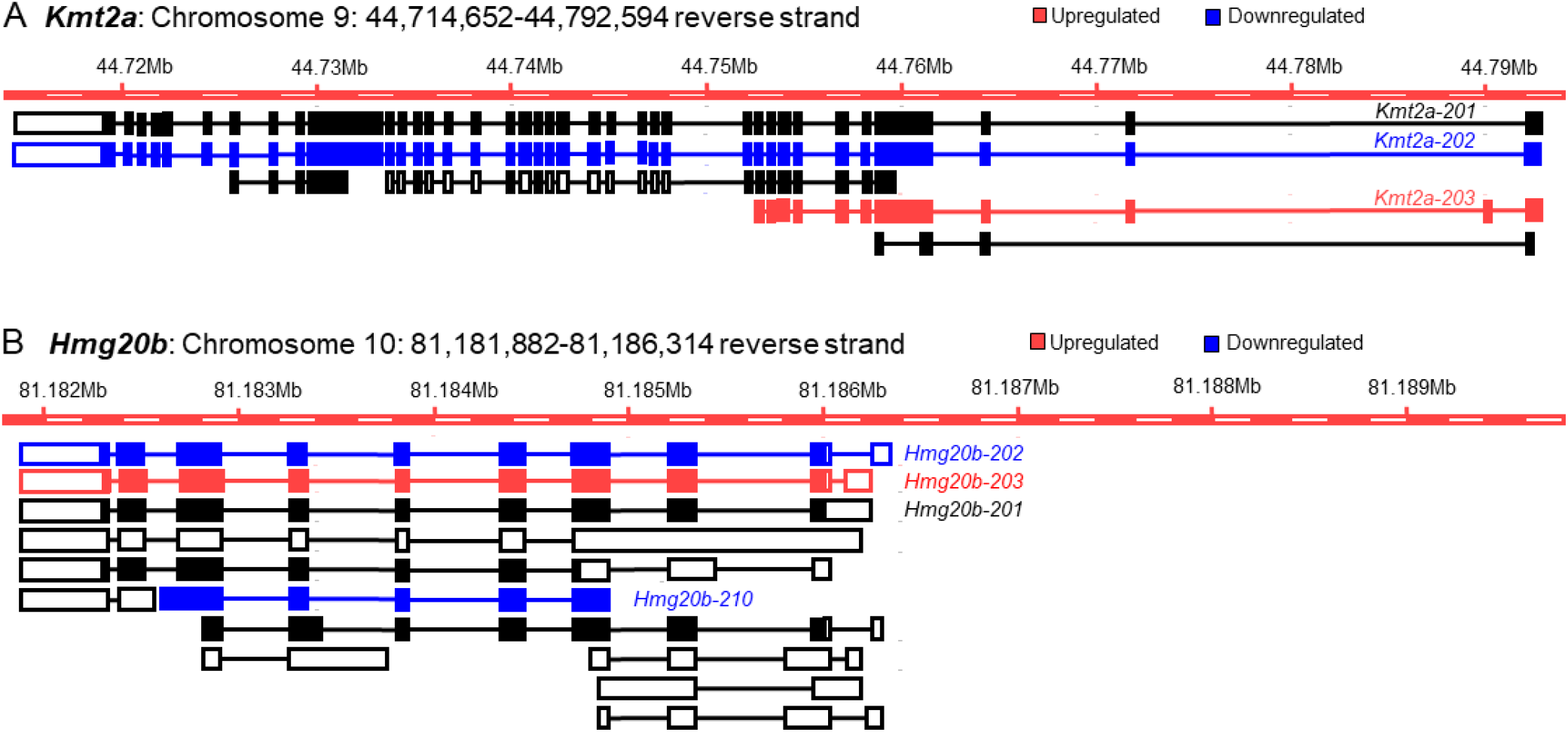
Impact of differentially expressed transcript variants on the upregulated genes. GE-based expression analyses identified both *Kmt2a* and *Hmg20b* as upregulated genes in mouse TS cells. We identified that transcript variant *Kmt2a-202* is significantly downregulated and expresses a full-length functional protein (A). However, this downregulation is masked by relatively more higher upregulation of another transcript variant of *Kmt2a* (*Kmt2a-203*), which expresses a truncated protein (**A**). We observed the downregulation of Hmg20b-202, which encodes a full-length protein, and Hmg20b-210, which encodes a truncated protein. Downregulation of the two transcript variants of *Hmg20b* remains unknown due to a higher upregulation of *Hmg20b*-*203* that encodes a full-length protein (**B**).

**Figure 7.**
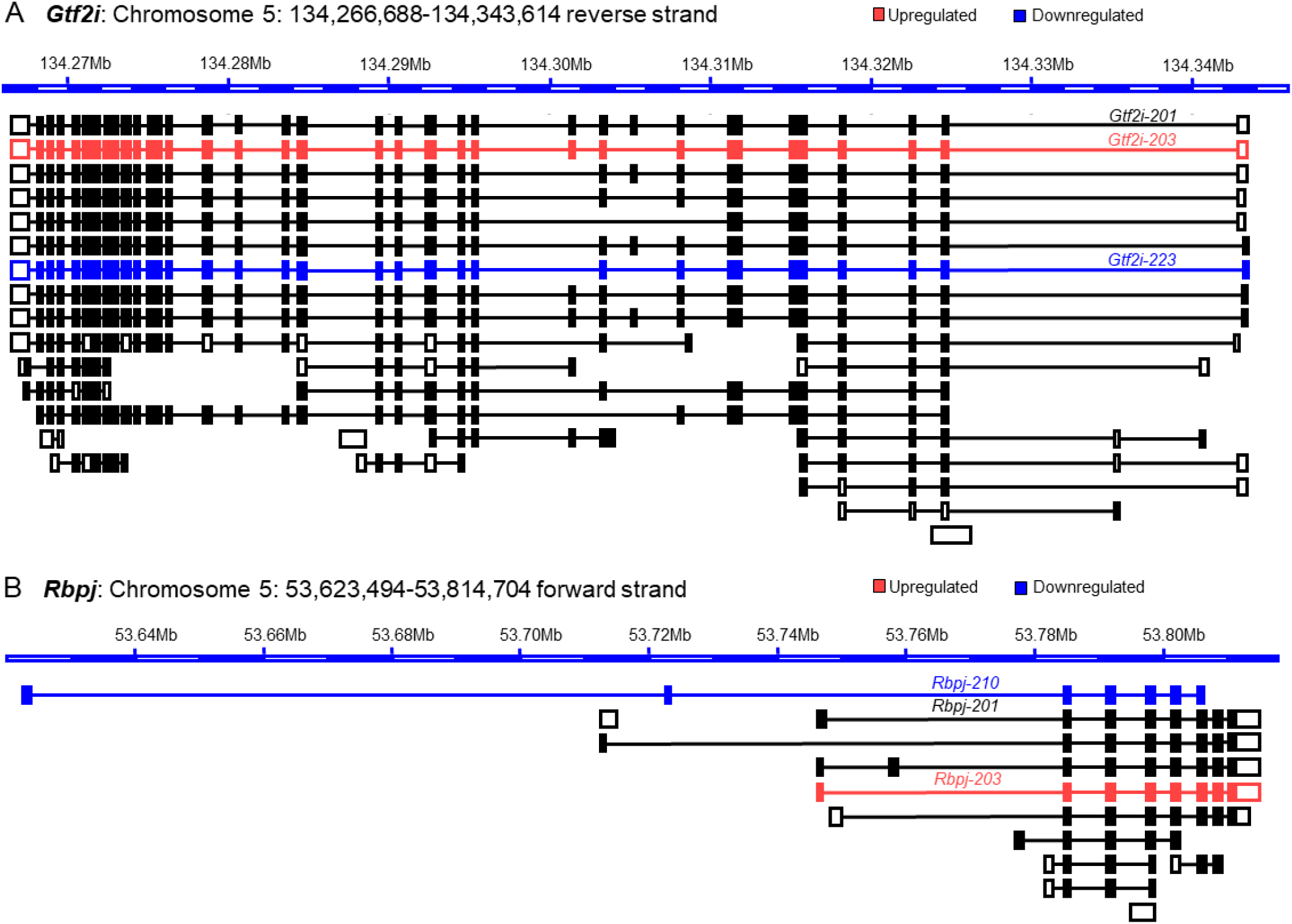
Impact of differential expression of transcript variants on the downregulated genes. GE-based expression analyses identified both *Gtf2i* and *Rbpj* as downregulated genes in mouse TS cells. We identified that transcript variant *Gtf2i-203* is significantly upregulated and lacks one exon (**A**). However, the upregulation of *Gtf2i-203* is masked by relatively higher downregulation of another transcript variant of *Gtf2i* (*Gtf2i-223*), which expresses a truncated protein lacking two proteincoding exons (**A**). We also observed that the upregulation of Rbpj-203, which encodes a full-length protein remains unknown due to a higher downregulation of another transcript variant of Rbpj (Rbpj-210) that encodes a protein with a different domain at either end (**B**).

**Figure 8.**
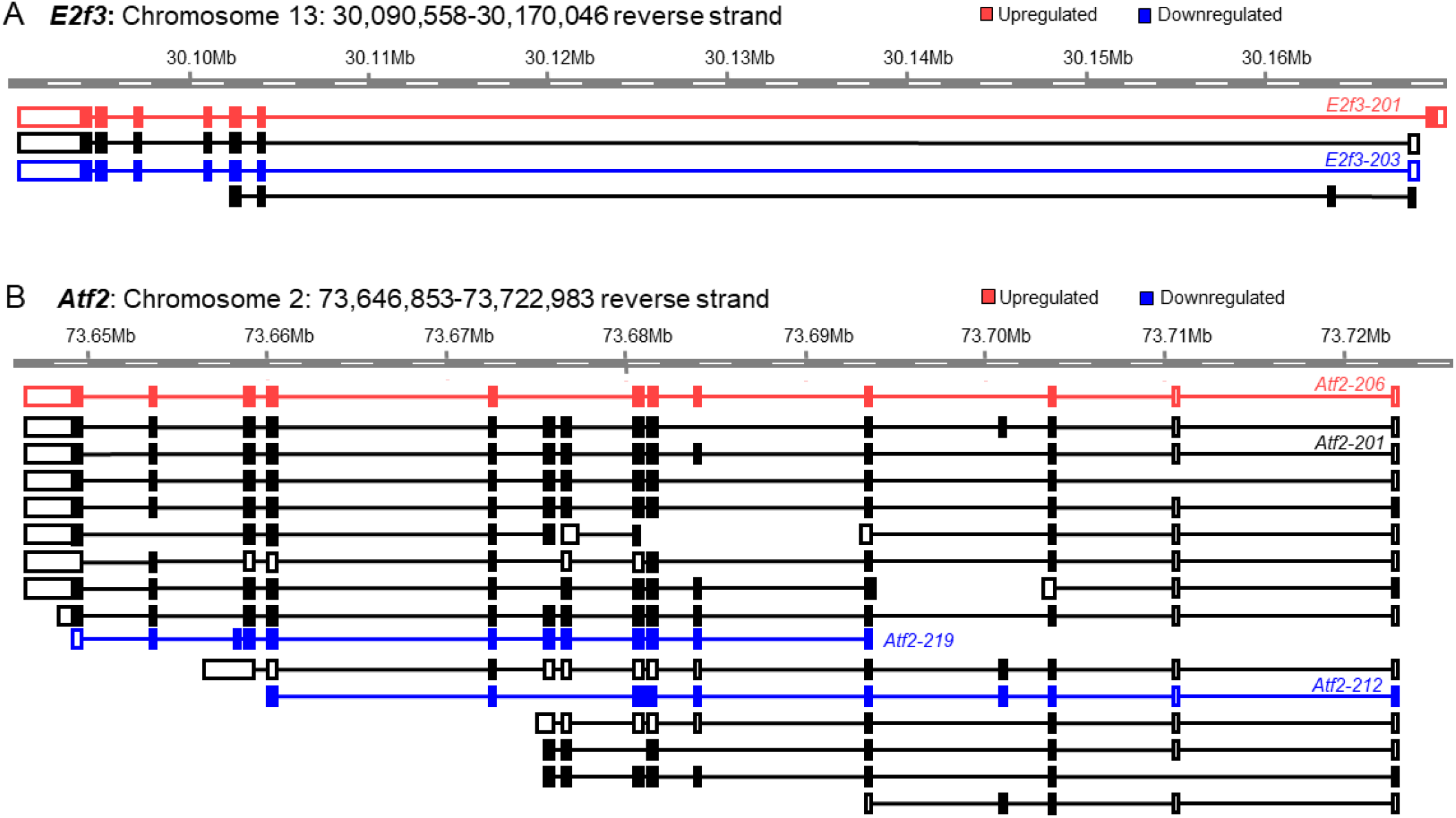
Impact of differential expression of transcript variants on the genes with insignificant differential expression. GE-based expression analyses did not identify both *E2f3* and *Atf2* as differentially expressed genes in mouse TS cells. However, we identified that a transcript variant of the *E2f3* gene (*E2f3-201*) was significantly upregulated, whereas another transcript variant of *E2f3* (*E2f3-203*) was significantly downregulated (**A**). Eventually, the upregulation of *E2f3-201* was masked by the downregulation of *E2f3-203*. Similarly, the significant upregulation of a transcript variant of *Atf2* (*Atf2-206*) and significant downregulation of two transcript variants of *Atf2* (*Atf2-219* and *Atf2-212*) remained undetected due to masking of *Atf2-206* results by those of *Atf2-219* and *Atf2212* (**B**).

A similar mechanism also underlies the masked upregulated transcript variants of *Gtf2i* (*Gtf2i-203*) due to the higher downregulation of *Gtf2i-223*, which does not encode 2 exons of the full-length protein (**Figure 7 A**). Another downregulated gene, *Rbpj*, also expresses an upregulated transcript variant, *Rbpj-203*, that expresses full-length protein, but this result remains unknown due to higher downregulation of another transcript variant, *Rbpj-210*, which encodes a protein truncated at the amino terminus and insertion at the carboxy terminus (**Figure 7 B**).

We also observed that both significantly upregulated and downregulated transcript variants may mask each other and their differential expressions remain unidentified during the analyses of GE-based gene expression (**Figure 8 A, B**). GE-based expression analyses did not identify both *E2f3* and *Atf2* as differentially expressed genes in mouse TS cells (**Figure 8 A, B**). We identified that a transcript variant of the *E2f3* gene (*E2f3-201*) was significantly upregulated. However, the upregulation of *E2f3-201* was masked by significant downregulation of another transcript variant of *E2f3* (*E2f3-203*)(**Figure 8 A**). Similarly, the significant upregulation of a transcript variant of *Atf2* (*Atf2-206*) remained masked by the significant downregulation of two transcript variants of *Atf2* (*Atf2-219* and *Atf2-212*) (**Figure 8B**).

## 4. Discussion

In RNA-Seq analyses, it is often assumed that any specific gene expresses only one transcript, leading to the inference that one gene encodes one mRNA and one protein. However, this assumption overlooks the fact that a single gene often can encode multiple transcripts due to alternative transcription start sites and alternative splicing^8, 32, 33^. These transcript variants can encode multiple in-frame peptides containing different structural and functional domains^34^. Moreover, some of the transcript variants can serve as noncoding regulatory RNAs or may undergo nonsense-mediated decay^35, 36^. Therefore, aggregating all transcript variants under a single gene name is not biologically accurate; future studies should include TE-based differential analyses of transcript variants.

This study was premised on the quantitative differences in the expression of the transcript variants of TF genes in mouse ES and TS cells. All the transcript variants are included under one gene name in GE-based analyses, but we have detected that the expression trends of these variants are not uniform. For instance, while a transcript variant expressed by a gene shows lineage-specific upregulation, another transcript expressed by the same gene can be downregulated simultaneously. These diverse patterns of the transcript variant expression in the same cell type may lead to erroneous estimations in GE-based differential gene expression analyses. Therefore, we quantified the potential errors arising from ignoring the differential expression of transcript variants (**Figures 4 and 5**).

Although the overall differential expression of TF genes and transcript variants were comparable between mouse ES and TS cells (**Figure 4**), deeper analyses depicted a different picture of discrepancy (**Figure 5-8**). We determined that only 14% (51 out of 365) of the upregulated genes did not express any downregulated or insignificant variants, and only 17% (48 out of 287) of the downregulated genes did not express any upregulated or insignificant transcript variants. Collectively, transcript variants expressed by more than 80% of the differentially expressed genes yielded inaccurate interpretations. Many upregulated genes included transcript variants that were either downregulated or had no significant changes (**Figures 4-8**). Many downregulated genes included upregulated transcript variants and transcripts without significant differences (**Figures 4-8**). Furthermore, genes that did not show significant changes in GE-based analyses expressed differentially expressed transcript variants (**Figures 4-8**).

More detailed analyses revealed that the transcript variants’ copy number (TPM value) plays an important role in determining the discrepancy between GE and TE-based analyses. Among the upregulated genes, approximately 86% of the low copy number transcript variants were discordant, whereas it was about 62% in the case of high copy number transcript (**Figure 5 G, H**). Among the downregulated genes, approximately 93% of the low copy number transcript variants were discordant, whereas it was about 60% in the case of high copy number transcripts (**Figure 5 G, H**). These observations indicate that low copy number transcript variants show a higher quantity of discrepancy. We suspect that statistical analyses sort the low copy number transcript variants more towards the insignificant group (**Figure 5G**). In contrast, the high copy number transcript variants stay more within the expected differentially expressed gene groups (**Figure 5H**). Remarkably, we detected a different pattern among the transcript variants corresponding to the insignificant group. While 90% of the low copy number transcript variants remained insignificant, it was reduced to 70% among the high copy number transcripts (**Figure 5 G, H**). The high copy number transcript variants were sorted from the insignificant to the differentially expressed group (**Figure 5 G, H**). Based on these findings, we can assume that if the sample size is increased, the genes or transcript variants in the insignificant group will decrease, and those in the upregulated or downregulated groups will increase. However, as most RNA-Seq studies include three samples in each group, we have used three libraries in each group.

We elucidated the underlying mechanism that results in the discrepancy between GE-based and TE-based differential expression analyses (**Figure 6-8**). A gene can be identified as upregulated on a GE basis if one or more of its transcript variants are highly upregulated despite one or more of its transcript variants’ low-level downregulations (**Figure 6**). Similarly, a gene can be identified as downregulated on a GE basis if one or more of its transcript variants are highly downregulated despite one or more of its transcript variants’ low-level upregulations (**Figure 7**). As expected, when a transcript variant of a gene shows upregulation and another variant shows downregulation, that may result in an insignificant difference in GE-based expression analysis despite the presence of two differentially expressed transcript variants (**Figure 8**).

If transcript variants are not included in RNA-seq analyses, we not only make errors, but we also disregard the roles of the transcript variants in a particular cell type. Cell lineage-specific transcriptional and posttranscriptional machinery generate the transcript variants of divergent molecular functions^37^. If we do not consider transcript variants, we also ignore transcriptional and posttranscriptional mechanisms. Transcript variants can also be expressed from alternative promoters, which suggests differential regulatory mechanisms involved in gene expression. Since alternative transcription start sites are not considered when the transcript variants are not analyzed, regulatory mechanisms involved in the initiation of transcription remain unknown. Genes also express transcript variants as long noncoding RNAs that have gene regulatory roles. Therefore, we also fail to uncover valuable gene regulatory information if we do not analyze the transcript variants.

## 5. Conclusions

During RNA-Seq analyses, GE values are considered, and TE values of the transcript variants are ignored to avoid relative procedural complexity. This study demonstrates the errors that are made when GE value-based differentially expressed genes are identified and transcript variants are not considered. Our results clearly indicate that gene expression analyses based on GE values are largely incorrect. RNA-Seq analyses should consider TE values of the transcript variants to identify their differential expression.

## Supplementary Materials

The following supporting information can be downloaded at: www.mdpi.com/xxx/s1, Figure S1: title; Table S1: title; Video S1: title.

## Author Contributions

M.A.K.R. conceptualization, supervision, funding acquisition, resources, and writing; K.V. data curation, methodology, investigation, formal analysis, and original draft preparation; R.M., Y.S. and A.M, software and data validation; P.E. F. review and editing. All authors have read and agreed on the contents of the manuscript.

## Institutional Review Board Statement

This study did not involve humans or animals.

## Informed Consent Statement

Not applicable

## Data Availability Statement

SRA, NCBI

## Acknowledgments

We acknowledge the editorial board of Cells to waive the publication fees. We also acknowledge Qiagen Bioinformatics for their continued support.

## Disclaimer/Publisher’s Note

The statements, opinions, and data contained in all publications are solely those of the individual author(s) and contributor(s) and not of MDPI and/or the editor(s). MDPI and/or the editor(s) disclaim responsibility for any injury to people or property resulting from any ideas, methods, instructions, or products referred to in the content.

## References

1. Zhang S, Pyne S, Pietrzak S, Halberg S, McCalla SG, Siahpirani AF, Sridharan R, Roy S. Inference of cell type-specific gene regulatory networks on cell lineages from single cell omic datasets. Nature Communications. 2023;14(1):3064. doi: 10.1038/s41467-023-38637-9.

2. Lowe R, Shirley N, Bleackley M, Dolan S, Shafee T. Transcriptomics technologies. PLoS Comput Biol. 2017;13(5):e1005457. Epub 20170518. doi: 10.1371/journal.pcbi.1005457. PubMed PMID: 28545146; PMCID: PMC5436640.

3. Chu Y, Corey DR. RNA sequencing: platform selection, experimental design, and data interpretation. Nucleic Acid Ther. 2012;22(4):271–4. Epub 20120725. doi: 10.1089/nat.2012.0367. PubMed PMID: 22830413; PMCID: PMC3426205.

4. Wang Z, Gerstein M, Snyder M. RNA-Seq: a revolutionary tool for transcriptomics. Nat Rev Genet. 2009;10(1):57–63. doi: 10.1038/nrg2484. PubMed PMID: 19015660; PMCID: PMC2949280.

5. Li W, Ballard J, Zhao Y, Long Q. Knowledge-guided learning methods for integrative analysis of multi-omics data. Comput Struct Biotechnol J. 2024;23:1945–50. Epub 20240430. doi: 10.1016/j.csbj.2024.04.053. PubMed PMID: 38736693; PMCID: PMC11087912.

6. Limbu MS, Xiong T, Wang S. A review of Ribosome profiling and tools used in Ribo-seq data analysis. Comput Struct Biotechnol J. 2024;23:1912–8. Epub 20240422. doi: 10.1016/j.csbj.2024.04.051. PubMed PMID: 38721586; PMCID: PMC11076270.

7. Samuels DS, Lybecker MC, Yang XF, Ouyang Z, Bourret TJ, Boyle WK, Stevenson B, Drecktrah D, Caimano MJ. Gene Regulation and Transcriptomics. Curr Issues Mol Biol. 2021;42:223–66. Epub 20201210. doi: 10.21775/cimb.042.223. PubMed PMID: 33300497; PMCID: PMC7946783.

8. Pal S, Gupta R, Kim H, Wickramasinghe P, Baubet V, Showe LC, Dahmane N, Davuluri RV. Alternative transcription exceeds alternative splicing in generating the transcriptome diversity of cerebellar development. Genome research. 2011;21(8):1260–72.

9. Reyes A, Huber W. Alternative start and termination sites of transcription drive most transcript isoform differences across human tissues. Nucleic acids research. 2018;46(2):582–92.

10. Alfonso-Gonzalez C, Hilgers V. (Alternative) transcription start sites as regulators of RNA processing. Trends in Cell Biology. 2024.

11. Xin D, Hu L, Kong X. Alternative promoters influence alternative splicing at the genomic level. PloS one. 2008;3(6):e2377.

12. Kelemen O, Convertini P, Zhang Z, Wen Y, Shen M, Falaleeva M, Stamm S. Function of alternative splicing. Gene. 2013;514(1):1–30. Epub 20120815. doi: 10.1016/j.gene.2012.07.083. PubMed PMID: 22909801; PMCID: PMC5632952.

13. Piazzi M, Bavelloni A, Salucci S, Faenza I, Blalock WL. Alternative splicing, RNA editing, and the current limits of next generation sequencing. Genes. 2023;14(7):1386.

14. Ha I, Roberts S, Maldonado E, Sun X, Kim LU, Green M, Reinberg D. Multiple functional domains of human transcription factor IIB: distinct interactions with two general transcription factors and RNA polymerase II. Genes Dev. 1993;7(6):1021–32. doi: 10.1101/gad.7.6.1021. PubMed PMID: 8504927.

15. Sonam D, Manoj BM. Non-coding transcript variants of protein-coding genes – what are they good for? RNA Biology. 2018;15(8):1025--31. doi: 10.1080/15476286.2018.1511675, note = PMID: 30146915.

16. Johnson KA, Krishnan A. Robust normalization and transformation techniques for constructing gene coexpression networks from RNA-seq data. Genome biology. 2022;23:1–26.

17. Conesa A, Madrigal P, Tarazona S, Gomez-Cabrero D, Cervera A, McPherson A, Szcześniak MW, Gaffney DJ, Elo LL, Zhang X. A survey of best practices for RNA-seq data analysis. Genome biology. 2016;17:1–19.

18. Jiang Z, Zhou X, Li R, Michal JJ, Zhang S, Dodson MV, Zhang Z, Harland RM. Whole transcriptome analysis with sequencing: methods, challenges and potential solutions. Cellular and molecular life sciences. 2015;72:3425–39.

19. Takahashi K, Yamanaka S. A decade of transcription factor-mediated reprogramming to pluripotency. Nature Reviews Molecular Cell Biology. 2016;17(3):183–93. doi: 10.1038/nrm.2016.8.

20. Kubaczka C, Senner CE, Cierlitza M, Araúzo-Bravo MJ, Kuckenberg P, Peitz M, Hemberger M, Schorle H. Direct Induction of Trophoblast Stem Cells from Murine Fibroblasts. Cell Stem Cell. 2015;17(5):557–68. Epub 20150924. doi: 10.1016/j.stem.2015.08.005. PubMed PMID: 26412560.

21. Johnston AD, Simões-Pires CA, Thompson TV, Suzuki M, Greally JM. Functional genetic variants can mediate their regulatory effects through alteration of transcription factor binding. Nature communications. 2019;10(1):3472.

22. Barrett T, Wilhite SE, Ledoux P, Evangelista C, Kim IF, Tomashevsky M, Marshall KA, Phillippy KH, Sherman PM, Holko M, Yefanov A, Lee H, Zhang N, Robertson CL, Serova N, Davis S, Soboleva A. NCBI GEO: archive for functional genomics data sets--update. Nucleic acids research. 2013;41(Database issue):D991-5. Epub 2012/11/30. doi: 10.1093/nar/gks1193. PubMed PMID: 23193258; PMCID: PMC3531084.

23. Tanaka S, Kunath T, Hadjantonakis AK, Nagy A, Rossant J. Promotion of trophoblast stem cell proliferation by FGF4. Science. 1998;282(5396):2072–5. Epub 1998/12/16. doi: 10.1126/science.282.5396.2072. PubMed PMID: 9851926.

24. Chakravarthi VP, Ratri A, Masumi S, Borosha S, Ghosh S, Christenson LK, Roby KF, Wolfe MW, Rumi MAK. Granulosa cell genes that regulate ovarian follicle development beyond the antral stage: The role of estrogen receptor β. Mol Cell Endocrinol. 2021;528:111212. Epub 2021/03/08. doi: 10.1016/j.mce.2021.111212. PubMed PMID: 33676987.

25. Khristi V, Chakravarthi VP, Singh P, Ghosh S, Pramanik A, Ratri A, Borosha S, Roby KF, Wolfe MW, Rumi MAK. ESR2 regulates granulosa cell genes essential for follicle maturation and ovulation. Mol Cell Endocrinol. 2018;474:214–26. Epub 2018/03/28. doi: 10.1016/j.mce.2018.03.012. PubMed PMID: 29580824.

26. Khristi V, Ratri A, Ghosh S, Pathak D, Borosha S, Dai E, Roy R, Chakravarthi VP, Wolfe MW, Karim Rumi MA. Disruption of ESR1 alters the expression of genes regulating hepatic lipid and carbohydrate metabolism in male rats. Mol Cell Endocrinol. 2019;490:47–56. Epub 2019/04/12. doi: 10.1016/j.mce.2019.04.005. PubMed PMID: 30974146.

27. Lambert SA, Jolma A, Campitelli LF, Das PK, Yin Y, Albu M, Chen X, Taipale J, Hughes TR, Weirauch MT. The Human Transcription Factors. Cell. 2018;172(4):650–65. doi: 10.1016/j.cell.2018.01.029. PubMed PMID: 29425488.

28. Nelder JA, Wedderburn RW. Generalized linear models. Journal of the Royal Statistical Society Series A: Statistics in Society. 1972;135(3):370–84.

29. Lin J, Khan M, Zapiec B, Mombaerts P. Efficient derivation of extraembryonic endoderm stem cell lines from mouse postimplantation embryos. Scientific reports. 2016;6:39457. Epub 2016/12/20. doi: 10.1038/srep39457. PubMed PMID: 27991575; PMCID: PMC5171707.

30. Ralston A, Cox BJ, Nishioka N, Sasaki H, Chea E, Rugg-Gunn P, Guo G, Robson P, Draper JS, Rossant J. Gata3 regulates trophoblast development downstream of Tead4 and in parallel to Cdx2. Development. 2010;137(3):395–403. doi: 10.1242/dev.038828. PubMed PMID: 20081188.

31. Takahashi K, Yamanaka S. Induction of pluripotent stem cells from mouse embryonic and adult fibroblast cultures by defined factors. Cell. 2006;126(4):663–76. Epub 2006/08/15. doi: 10.1016/j.cell.2006.07.024. PubMed PMID: 16904174.

32. Stamm S, Ben-Ari S, Rafalska I, Tang Y, Zhang Z, Toiber D, Thanaraj T, Soreq H. Function of alternative splicing. Gene. 2005;344:1–20.

33. Ashkenas J. Gene regulation by mRNA editing. American journal of human genetics. 1997;60(2):278.

34. Ray TA, Cochran K, Kozlowski C, Wang J, Alexander G, Cady MA, Spencer WJ, Ruzycki PA, Clark BS, Laeremans A. Comprehensive identification of mRNA isoforms reveals the diversity of neural cell-surface molecules with roles in retinal development and disease. Nature communications. 2020;11(1):3328.

35. Sun B, Chen L. Mapping genetic variants for nonsense-mediated mRNA decay regulation across human tissues. Genome Biology. 2023;24(1):164.

36. Marchese FP, Raimondi I, Huarte M. The multidimensional mechanisms of long noncoding RNA function. Genome biology. 2017;18:1–13.

37. Ju W, Greene CS, Eichinger F, Nair V, Hodgin JB, Bitzer M, Lee Y-s, Zhu Q, Kehata M, Li M. Defining cell-type specificity at the transcriptional level in human disease. Genome research. 2013;23(11):1862–73.

